# Examining the Neural Correlates of Error Awareness in a Large fMRI Study

**DOI:** 10.1101/2022.01.06.475224

**Authors:** Gezelle Dali, Méadhbh Brosnan, Jeggan Tiego, Beth P. Johnson, Alex Fornito, Mark A. Bellgrove, Robert Hester

## Abstract

Goal-directed behaviour is dependent upon the ability to detect errors and implement appropriate post-error adjustments. Accordingly, several studies have explored the neural activity underlying error-monitoring processes, identifying the insula cortex as crucial for error awareness and reporting mixed findings with respect to the anterior cingulate cortex. Variable patterns of activation have previously been attributed to insufficient statistical power. We therefore sought to clarify the neural correlates of error awareness in a large event-related functional magnetic resonance imaging (MRI) study. Four hundred and two healthy participants undertook the error awareness task, a motor Go/No-Go response inhibition paradigm in which participants were required to indicate their awareness of commission errors. Compared to unaware errors, aware errors were accompanied by significantly greater activity in a network of regions including the insula cortex, supramarginal gyrus, and midline structures such as the anterior cingulate cortex and supplementary motor area. Error awareness activity was related to indices of task performance and dimensional measures of psychopathology in selected regions including the insula, supramarginal gyrus and supplementary motor area. Taken together, we identified a robust and reliable neural network associated with error awareness.

## Introduction

Error processing facilitates goal-directed behaviour through error detection and the execution of appropriate post-error adjustments. Within error processing, it is possible to delineate between errors made with and without conscious recognition. Although error processing can proceed in the absence of awareness, conscious perception of errors may subserve the implementation of remedial behaviours. Critically, deficient error awareness has been associated with symptoms of inattention, lack of insight and perseverative behaviour in several clinical conditions such as attention-deficit hyperactivity disorder (ADHD; O’Connell et al. 2009), autism spectrum disorder (ASD; Klein et al. 2013b) and substance use disorder (Hester et al, 2009), providing impetus for investigating the underlying neurobiology of error awareness.

In performance monitoring tasks, errors are largely associated with an event-related potential (ERP) signature comprising the error-related negativity (ERN) and the error positivity (Pe). The ERN is a negative deflection with a fronto-central distribution that appears 50-100ms following an error (Gehring et al. 1993), whereas the Pe is a positive deflection with an approximate latency of 300-500ms occurring over a centro-parietal location (Falkenstein et al. 1991). Neuroimaging and source localisation studies have identified the anterior cingulate cortex (ACC) as the source of the ERN (Debener et al. 2005; van Veen and Carter 2006). Indeed, the ACC is consistently implicated in performance monitoring tasks and is suggested to navigate the selection and evaluation of goal-directed behaviours (Holroyd and Yeung 2012). The source of the Pe, however, remains somewhat equivocal, with reports that it arises from activity in the rostral ACC (rACC; Herrmann et al. 2004; Van Boxtel et al. 2005), prefrontal (Masina et al. 2019) and parietal cortices (van Veen and Carter 2006; O’Connell et al. 2007).

With regard to error awareness, most electrophysiological studies argue that the ERN is unaffected by conscious error perception. This pattern has been observed in a myriad of tasks, including anti-saccade tasks (Nieuwenhuis et al. 2001; Endrass et al. 2007), Go/No-Go error awareness tasks (O’Connell et al. 2007; Dhar et al. 2011) and visual discrimination tasks (Steinhauser and Yeung 2010; Endrass et al. 2012). Contrastingly, the Pe has been found to be selectively modulated by error awareness such that greater amplitudes are observed following aware errors (Nieuwenhuis et al. 2001; O’Connell et al. 2007; Steinhauser and Yeung 2010; Dhar et al. 2011; Hoffmann and Beste 2015). There are, however, studies which have demonstrated that both the ERN and Pe are sensitive to error awareness (Scheffers and Coles 2000; Maier et al. 2008; Hewig et al. 2011; Wessel et al. 2011; Shalgi and Deouell 2012). Given such findings, the ERN has been proposed to be the foremost indication that an error has occurred and may serve as a feedforward input signal into systems that are more responsible for error awareness (Murphy et al. 2012; Wessel 2012), whereas the Pe reflects the accumulation of information that leads to error awareness (Klein et al. 2013b).

Neuroimaging studies on error awareness have been found to accord with electrophysiological findings. Consistent with findings on the Pe, a network of frontal and parietal regions has been implicated in error awareness, namely the bilateral inferior parietal and bilateral mid-frontal cortices (Hester et al. 2005; Harsay et al. 2012; Orr and Hester 2012). The insula cortex – largely the anterior insula cortex (AIC) – is also widely recognised to be selectively modulated by error awareness (Klein et al. 2013b). While the insula is unlikely to generate the Pe directly, it has been suggested to indirectly elicit the Pe through its functional connections with frontal and parietal cortices (Klein et al. 2007a). Corroborating findings on the ERN, the relationship between awareness and the ACC remains a topic of contention. Several earlier studies that have found ACC activity to be greater for errors than correct responses have discerned no difference in activity between aware and unaware errors (e.g., Hester et al. 2005; Klein et al. 2007a). In contrast to the majority of earlier studies, recent investigations have reported dorsal ACC (dACC) sensitivity to error awareness, with increased activity observed during aware errors (Harsay et al. 2012; Harsay et al. 2018).

Although heterogeneity in imaging modalities, sample characteristics and study designs may contribute to discordant neuroimaging findings, they are unlikely to explain variation observed across several error awareness studies (Wessel 2012). Instead, disparities in distinguishing ACC activity patterns between aware and unaware errors may be attributed to inadequate statistical power associated with small sample sizes. For example, we have previously found no difference in ACC activity between aware and unaware errors in samples of 13 (Hester et al. 2005) and 16 (Hester et al. 2009a) participants, however have found greater dACC activity for aware errors in a sample of 27 participants (Hester et al. 2012). Importantly, when the samples of these three studies were collated, a significant effect of awareness on dACC activity was observed (Orr and Hester 2012). Insufficient statistical power thus seems a robust explanation for these mixed neuroimaging findings (Button et al. 2013; Poldrack et al. 2017). Indeed, low power is a pertinent problem for task-based neuroimaging studies where there are often a small number of observations and few participants (Cremers et al. 2017; Turner et al. 2018). Although recent work has begun to address the reproducibility of brain imaging (Bossier et al. 2020), relatively few functional imaging replication studies have been conducted in this area of research. In light of this shortcoming, the neural correlates of error awareness and the influence of measures of task and individual differences warrants further examination.

Here, we set out to confirm previous investigations using the motor Go/No-Go error awareness task (Hester et al. 2005), in a large, community-based sample. Behavioural performance on the error awareness task and corresponding event-related neuroimaging were used to assess the neural mechanisms associated with error awareness. Based on the reviewed literature, we hypothesise that aware errors will be accompanied by greater activity in a network of regions including the insula, parietal and mid-frontal cortices, and midline structures such as the ACC. Further, we extend upon previous investigations by exploring whether existing findings from clinical samples (e.g., ADHD, ASD) are also apparent in larger-scale healthy samples. Specifically, we examined whether variance in awareness-related neural activity is accounted for by individual differences in dimensional measures of psychopathology including ADHD, ASD, and impulsivity.

## Materials and Methods

### Participants

Participants were recruited via Monash University Clayton campus, social media and newspaper advertisements along with experimenter networks. All participants were right-handed and had normal or corrected-to-normal vision. Participants were excluded if they were colour blind or reported any history of neurological or psychiatric illness, including head injury, previous usage of psychotropic medication or substance use disorder. All participants provided written informed consent and were reimbursed for participation. The study received approval by the Monash University Human Research Ethics Committee for meeting the research standards prescribed by the National Health and Medical Research Council (CF12/3072 – 2012001562).

Four hundred and seventy-three participants completed the event-related fMRI protocol. Participants were subsequently excluded due to missing functional runs (*n* = 4) or behavioural data (*n* = 22), no signalling of aware errors (*n* = 37), corrupted functional data (*n* = 6), or distorted anatomical data (*n* = 2). The final sample with complete behavioural and neuroimaging data comprised 402 participants (female, 54.22%; *M*_*age*_ = 23.64 years, *SD =* 5.45; age range: 18-50 years). Of those, 20 participants (female, 50%; *M*_*age*_ *=* 25.55 years, *SD* = 7.25) did not have questionnaire data available. Further information on participant age can be found in Table 1A of the Supplementary Material.

**Table 1.** Behavioural Performance: Inhibition Accuracy, Error Awareness and Reaction Time on the Error Awareness Task

| Category | Mean (SD) |
| --- | --- |
| Inhibition accuracy % | 53.59 (19.31) |
| Total errors | 66.45 (29.05) |
| Error awareness % | 86.47 (11.74) |
| Reaction time (ms) |  |
| Go trial | 517.66 (80.60) |
| Aware trial | 489.14 (107.28) |
| Unaware trial | 507.37 (117.33) |
| Post-No-Go reaction time adjustment (ms)<sup>a</sup> |  |
| Aware error | +5.94 (49.66) |
| Unaware error | +19.22 (93.21) |
| Correct inhibition | +7.89 (44.24) |

<sup>a</sup>Post-No-Go reaction time – pre-No-Go reaction time. Post-No-Go reaction time is taken from the Go trial that succeeds the No-Go error by three trials.

### Experimental Design

Participants were administered a battery of self-report measures designed to assess a comprehensive range of psychopathological characteristics. The battery comprised the Barratt Impulsiveness Scale, Version 11 (BIS-11; Barratt and Patton 1983) to assess impulsivity, Conners’ Adult ADHD Rating Scales – Self Report: Long Version (CAARS - S:L; Conners 1998) to assess ADHD-like behaviours, the Behavioural Inhibition/Activation Systems Scale (BIS/BAS; Carver and White 1994) to measure sensitivity to avoidance and approach motivation, the Autism-Spectrum Quotient (AQ; Baron-Cohen et al. 2001) to assess autistic traits, and the Hospital Anxiety and Depression Scale (HADS; Zigmond and Snaith 1983) to assess anxiety and depression traits.

### Error Awareness Task

The error awareness task (see Figure 1) is a Go/No-Go motor inhibition task that presents a serial stream of colour words in a congruent or incongruent colour. Previously, we employed an awareness task with two competing inhibition rules (a repeat No-Go rule and colour No-Go rule). To address concerns that introducing two No-Go rules potentially contaminates the BOLD signal, we opted to remove the repeat No-Go rule. Pilot testing confirmed that unaware error rates with a single No-Go rule were consistent with our previous work (see Table 2A of the Supplementary Material for a summary of pilot data). Thus, participants were required to respond to the incongruent trials (Go trials) with a left button press, while withholding their response when the word and colour were congruent (No-Go trials). To indicate error awareness, participants were trained to forego making a standard ‘Go’ response and instead execute a right button press following any commission error. Erroneous No-Go trials were those in which a participant failed to withhold a response. To classify erroneous trials for data analysis, unaware errors were those in which the participant responded with a left button press on the No-Go trial and again on the following Go trial. Any deviation from this pattern of response on a No-Go trial or following a No-Go error was classified as an aware error (Figure 1).

**Table 2.**
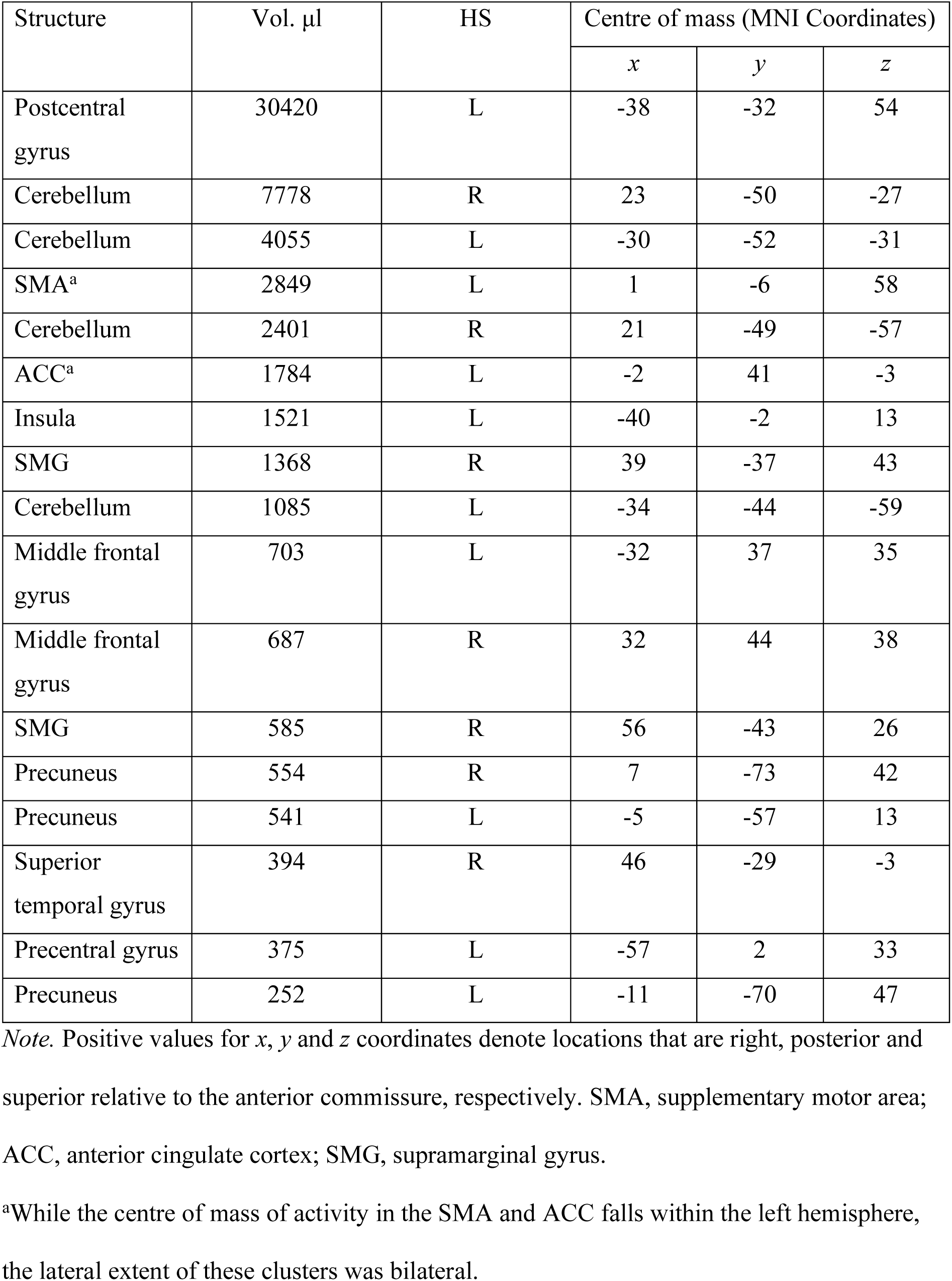
Regions that Showed Significantly Greater BOLD Signal for Aware Errors than Unaware Errors

| Structure | Vol. μl | HS | Centre of mass (MNI Coordinates) | | |
| --- | --- | --- | --- | --- | --- |
|  |  |  | *x* | *y* | *z* |
| Postcentral gyrus | 30420 | L | -38 | -32 | 54 |
| Cerebellum | 7778 | R | 23 | -50 | -27 |
| Cerebellum | 4055 | L | -30 | -52 | -31 |
| SMA<sup>a</sup> | 2849 | L | 1 | -6 | 58 |
| Cerebellum | 2401 | R | 21 | -49 | -57 |
| ACC<sup>a</sup> | 1784 | L | -2 | 41 | -3 |
| Insula | 1521 | L | -40 | -2 | 13 |
| SMG | 1368 | R | 39 | -37 | 43 |
| Cerebellum | 1085 | L | -34 | -44 | -59 |
| Middle frontal gyrus | 703 | L | -32 | 37 | 35 |
| Middle frontal gyrus | 687 | R | 32 | 44 | 38 |
| SMG | 585 | R | 56 | -43 | 26 |
| Precuneus | 554 | R | 7 | -73 | 42 |
| Precuneus | 541 | L | -5 | -57 | 13 |
| Superior temporal gyrus | 394 | R | 46 | -29 | -3 |
| Precentral gyrus | 375 | L | -57 | 2 | 33 |
| Precuneus | 252 | L | -11 | -70 | 47 |

*Note.* Positive values for *x*, *y* and *z* coordinates denote locations that are right, posterior and superior relative to the anterior commissure, respectively. SMA, supplementary motor area; ACC, anterior cingulate cortex; SMG, supramarginal gyrus.
<sup>a</sup>While the centre of mass of activity in the SMA and ACC falls within the left hemisphere, the lateral extent of these clusters was bilateral.

**Figure 1.**
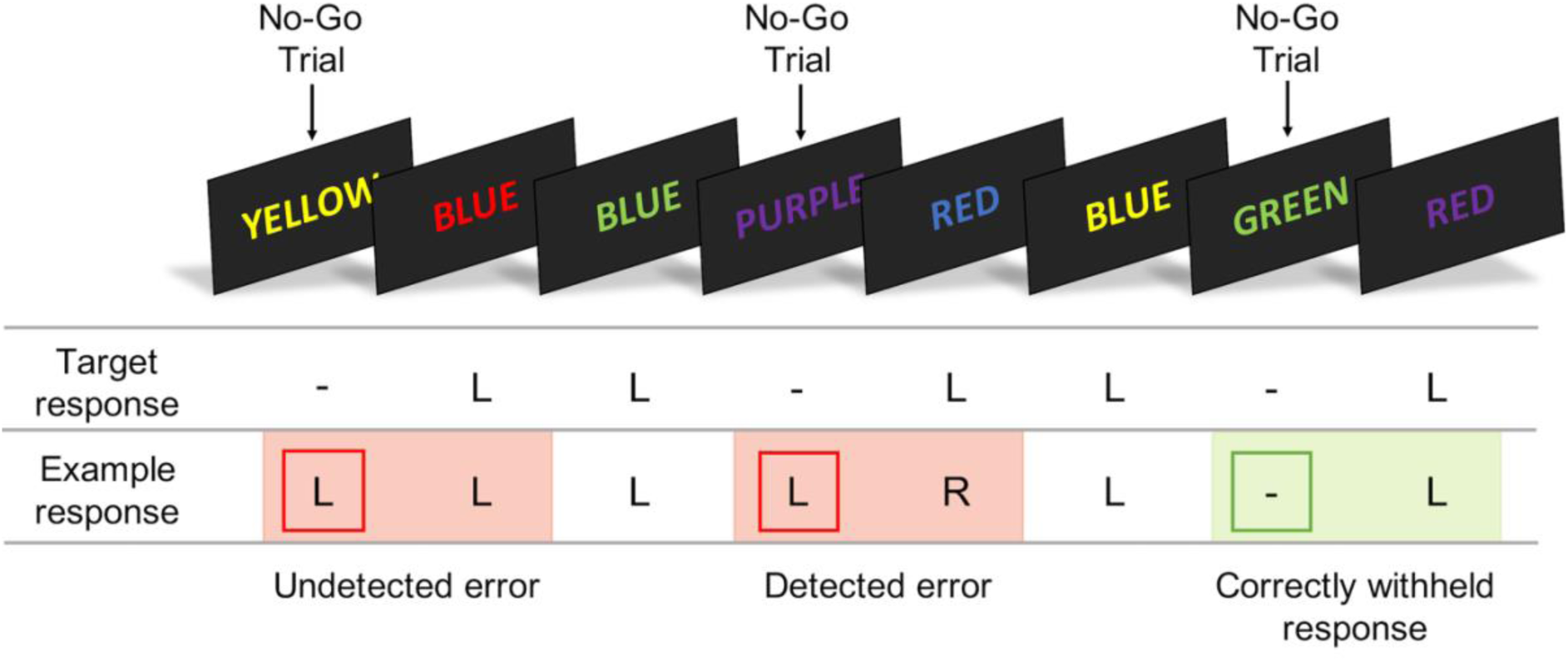
*Note*. The error awareness task presents a serial stream of colour words, in a congruent (No-Go trial) or incongruent (Go trial) colour. Participants respond to Go trials using a left button press (‘L’) while withholding their response to No-Go trials. To indicate error awareness, participants forgo making a standard ‘Go’ response and instead execute a right button press (‘R’) on the trial following the commission error. The task comprised six blocks, each with 175 trials. Across all blocks, participants were administered 900 Go trials and 150 No-Go trials. All stimuli were presented for 900 ms followed by a 600 ms inter-stimulus interval.

The task comprised six blocks, each with 175 trials. Across all blocks, participants were administered 900 Go trials and 150 No-Go trials. All stimuli were presented for 900 ms followed by a 600 ms inter-stimulus interval. An event-related design was employed, distributing the No-Go trials pseudo-randomly throughout the serial presentation of Go trials. Events of interest were adequately separated in order to analyse correct and failed response inhibition events separately without cross-contamination. The number of Go trials separating No-Go trials ranged between 1 and 12 (*M =* 6.23; *SD* = 2.55).

### Image Acquisition

Scanning was conducted between August, 2013 and July, 2017, at Monash Biomedical Imaging (Victoria, Australia). Images were acquired using a Siemens Skyra 3-Tesla MRI scanner with a 32-channel head coil. High resolution T1-weighted structural MPRAGE images (TE = 2.07 ms, TR = 2300 ms, FOV = 256 mm, flip angle = 9 degrees, thickness = 1 mm isotropic, sagittal slices) were acquired prior to functional imaging to enable activation localisation and for spatial normalisation. Functional images were acquired using a gradient-echo pulse (EPI) sequence (TE = 30 ms, TR = 2460 ms, FOV = 190 mm, flip angle = 90 degrees, 44 contiguous transversal slices of 3.0 mm thickness). The error awareness task was presented using E-Prime software (version 2.0; Psychology Software Tools) on a Cambridge 32-inch BOLD screen which was reflected onto a mirror visor positioned in the radio frequency head coil. Participants responded to each stimulus using their right hand, entering their responses using two buttons on a four-button MR-compatible response pad (Fiber-Optic response pads; Current Designs).

### Statistical Analysis

#### Behavioural Analysis

Behavioural data analyses were undertaken in the programming language R using the *stats* package (R Core Team 2017), with the addition of the *psych* (Revelle 2020), *afex* (Singmann et al. 2020) and *emmeans* (Russell et al. 2020) packages. Effect sizes were calculated using the *effectsize* package (Ben-Shachar et al. 2020). Assumptions were tested, and non-parametric analyses were computed under violations of normality. Greenhouse-Geisser-adjusted degrees of freedom and *p-*values are reported under violations of sphericity. Post-hoc tests were undertaken using Tukey’s method for multiple comparisons. *P*-values were otherwise adjusted using Holm procedures. Alpha was set to .05 for all analyses. The number of trials available for our behavioural analyses are outlined in Table 3A of the Supplementary Material. The full reproducible code for the current results has been made publicly available online (https://osf.io/hrba7/).

**Table 3.** Lasso Regression Coefficients

|  | Outcome |  |  |  |  |  |  |  |  |  |
| --- | --- | --- | --- | --- | --- | --- | --- | --- | --- | --- |
|  | Error awareness |  | L-insula |  | ACC |  | R-SMG |  | R-Middle frontal gyrus |  |
| Predictor | Lasso coeff. | non-zero (%) | Lasso coeff. | non-zero (%) | Lasso coeff. | non-zero (%) | Lasso coeff. | non-zero (%) | Lasso coeff. | non-zero (%) |
| BIS-11 – attentional | . | 56 | . | 56 | . | 26 | -0.05 | 58 | . | 33 |
| BIS-11 – motor | -0.07 | 86 | . | 46 | . | 30 | 0.11 | 78 | 0.05 | 74 |
| BIS-11 – non-planning | -0.03 | 90 | . | 38 | . | 48 | -0.01 | 52 | . | 34 |
| AQ – social skill | . | 48 | . | 40 | . | 56 | -0.15 | 82 | . | 46 |
| AQ – attention switching | . | 68 | . | 50 | . | 22 | . | 54 | . | 37 |
| AQ – attention to detail | . | 64 | .0003 | 60 | . | 44 | 0.04 | 64 | . | 47 |
| AQ – communication | . | 58 | . | 56 | . | 20 | 0.16 | 92 | 0.05 | 71 |
| AQ – imagination | . | 54 | 0.02 | 76 | . | 34 | 0.03 | 64 | 0.13 | 92 |
| BIS/BAS – BAS drive | . | 68 | . | 56 | . | 30 | 0.04 | 66 | 0.05 | 72 |
| BIS/BAS – BAS fun | . | 50 | . | 42 | . | 24 | -0.12 | 74 | . | 25 |
| BIS/BAS – BAS reward | . | 62 | . | 42 | . | 34 | 0.01 | 54 | . | 41 |
| BIS/BIS – BIS score | 0.04 | 90 | . | 68 | . | 30 | 0.03 | 60 | 0.01 | 53 |
| HADS – anxiety | . | 52 | . | 50 | . | 30 | 0.02 | 64 | . | 49 |
| HADS – depression | . | 52 | 0.10 | 90 | . | 34 | 0.19 | 98 | . | 79 |
| CAARS – attention | . | 42 | . | 38 | . | 22 | . | 38 | . | 43 |
| CAARS – hyperactivity | . | 46 | . | 52 | . | 24 | 0.11 | 54 | . | 25 |
| CAARS – impulsivity | . | 48 | . | 48 | . | 22 | -0.04 | 60 | . | 37 |
| CAARS – self-concept | . | 46 | . | 42 | . | 54 | . | 36 | . | 32 |
| CAARS – DSM attention | . | 34 | . | 50 | . | 20 | . | 42 | -0.05 | 49 |
| CAARS – DSM hyperactivity | . | 38 | . | 40 | . | 22 | -0.14 | 60 | . | 23 |
| CAARS – DSM ADHD | . | 6 | . | 4 | . | 6 | -0.01 | 20 | . | 24 |
| CAARS – index | . | 64 | . | 18 | . | 12 | -0.02 | 40 | . | 24 |

*Note.* Results are presented as standardised regression coefficients. To determine the robustness of the variable selection, each lasso model was computed on 500 bootstrap samples. The percentage of non-zero bootstrap samples is reported for each variable alongside the coefficients. BIS, Barratt Impulsiveness Scale, Version 11; AQ, Autism Spectrum Quotient; BIS/BAS, Behavioural Inhibition/Avoidance Scale; HADS, Hospital Anxiety and Depression Scale; CAARS, Conners' Adult ADHD Rating Scales; ACC, anterior cingulate cortex; SMG, supramarginal gyrus.

The error awareness task is not optimised to analyse response speed adjustments following errors as participants are required to make an awareness button press on the first post-error trial. Switching to the awareness button typically results in abnormally fast reaction times on the Go trials following the error. Response speed adjustments following No-Go trials were therefore determined by calculating the difference in reaction time for the Go trial following the No-Go trial by three trials and the Go trial immediately preceding the No-Go trial (a subtraction of the pre-error Go reaction time from the third Go reaction time after the No-Go trial).

#### Neuroimaging Analysis

Neuroimaging analyses were undertaken using AFNI software (Cox 1996). Data analysis procedures followed those implemented in studies with similar experimental paradigms (e.g., Hester et al. 2012). Behavioural data were used to categorise trial events into the following regressors: correct inhibitions, unaware errors and aware errors. Activation outside of the brain was removed using edge detection techniques. Following image reconstruction, the time series data were time shifted (using Fourier interpolation) to remove differences in slice acquisition times and then motion corrected using 3D volume registration (least-squares alignment of three translational and three rotational parameters).

Using the BLOCK basis function, separate haemodynamic impulse response functions (IRFs) were computed at 2.46 s temporal resolution for aware errors, unaware errors and correct inhibitions. To avoid confounding the baseline and event-related activity estimates, rest and omission errors were included as regressors of no interest. A multiple regression program (3dDeconvolve) determined the best fitting gamma variate function for these IRFs. The area under the curve of the gamma variate function was expressed as a percentage of the area under the baseline. The baseline in this design refers to task-related Go-trial processing that remains once the variance of the other events has been removed. The percentage area (event-related activation) map voxels were re-sampled at 1 mm resolution, then spatially normalised to standard MNI space and spatially blurred with a 3 mm isotropic root mean squared Gaussian kernel.

Group activation maps were obtained using a paired samples *t-*test (3dttest++) against the null hypothesis of no event-related activation differences between aware and unaware errors. Significant voxels passed a voxel-wise statistical threshold (*t* = 6.60, *p* = 1.0 × 10^−10^) and were required to be part of a 250 μl cluster of significant contiguous voxels. This method of combining probability and cluster thresholding sought to maximise power while minimising the likelihood of false-positives. ANFI’s 3dClustSim was provided with the number of voxels in the group map, the spatial correlation of the voxels, and the voxel-wise threshold. A series of Monte Carlo simulations (10,000 iterations) were then undertaken to determine the frequency of varying sized clusters produced by chance. From this frequency distribution, we selected the cluster size that occurred less than 1% of the time by chance, to provide a threshold of *p* = .010, corrected. Using this method for the current sample resulted in a highly liberal cluster-wise threshold (< 1 μl). We thus opted for a cluster-wise threshold of 250 μl as it is far more conservative and is moreover comparable with previous studies (e.g., Hester et al. 2005). Mean activity estimates for each event were derived for clusters in the whole brain map using the program 3DRoiStats. The estimates were used in assessing the relationship between neural activity and measures of task performance and individual differences.

#### Linking Neural Activity to Psychopathological Traits

Lasso (least absolute shrinkage and selection operator) regression was employed to determine a subset of the dimensional psychopathological measures that best predict error awareness and mean activity estimates for the insula cortex, anterior cingulate cortex (ACC), supramarginal gyrus (SMG) and middle frontal gyrus. Lasso is a modified form of least squares regression that applies a regularisation parameter (λ) to determine the variables that best predict the outcome measure (Tibshirani 1996). The regularisation parameter shrinks coefficients to zero for irrelevant covariates in order to minimise prediction error and reduce overfitting. The optimal penalty term was determined using a 10-fold cross-validation. By enforcing sparsity, lasso regression provides a principled way of identifying a subset of predictors that have the strongest influence on the dependent variable (Tibshirani 1996). Lasso generalised linear models were computed in the programming language R (R Core Team 2017) using the *glmnet* package (Friedman et al. 2010). The main independent variables were subscale scores from each of the aforementioned psychopathological questionnaires. Although a whole-brain approach was used to explore the regions associated with awareness, a more focused subset of areas was selected as dependent variables to investigate how awareness activity is related to psychopathological traits. Dependent variables were therefore error awareness, and mean aware activity estimates for four clusters identified in our imaging analysis (insula cortex, ACC, SMG and middle frontal gyrus). These clusters were selected due to theoretical relevance and previous findings of sensitivity to error awareness (Harsay et al. 2012; Orr and Hester 2012; Klein et al. 2013b). Five separate models were computed, one for each dependent variable. The analysis does not allow missing data, therefore cases with missing values were omitted. Little’s test indicated that data were missing completely at random, χ _2_(155) = 173.53, *p* = .155. All variables were standardised prior to analysis to generate *Z*-scores. A test statistic or *p-*value for lasso regression is still under development (Lockhart et al. 2014). Further, given the interest here is predictive performance and not statistical inference, results are presented as standardised regression weights alone. To determine the robustness of the variable selection, each lasso model was computed on 500 bootstrap samples. The percentage of non-zero bootstrap samples is reported for each variable alongside the coefficients.

## Results

### Behavioural Results

Performance indices are summarised in Table 1. Participants correctly withheld 53.59% of their responses on No-Go trials, and were aware of 86.47% of commission errors (error awareness range 21.74-99.11%). There was a non-significant weak association between awareness of errors and overall inhibition performance, *r*_*s*_ = -.10, *p =* .055. A repeated measures ANOVA revealed that the speed of response was significantly related to trial type, *F*(2, 798) = 21.37, *p* < .001, *η*_*p*_^2^ = .05. Post-hoc tests using the Tukey method for multiple comparisons indicated that reaction times were significantly faster for aware errors than for either unaware errors, *t*(798) = -4.22, *p* < .001, *d* = -.23, 95% CI [-.30, -.16], or correct Go responses, *t*(798) = -6.44, *p* < .001, *d* = -.15, 95% CI [-.22, -.08]. There was no significant difference in reaction time between correct responses and unaware errors, *t*(798) = 2.22, *p =* .069, *d* = -.08, 95% CI [-.15, -.01].

A repeated measures ANOVA was computed to compare reaction time adjustments across No-Go responses (correct inhibitions, aware error, unaware error). The results revealed an effect of No-Go response type on post-No-Go reaction time, *F*(1.52, 598.43) = 4.85, *p* = .015, *η*_*p*_ ^2^ = .01. Post-hoc tests indicated greater slowing of responses following unaware errors (+19ms) compared to aware errors (+5ms), *t*(786) = 2.91, *p =* .010, *d* = -.48, 95% CI [-.56, -.41], and correct No-Go responses (+7ms), *t*(786) = 2.41, *p =* .043, *d* = -.40, 95% CI [-.47, -.33]. There was no significant difference in post-No-Go reaction adjustments between correct responses and aware errors, *t*(786) = 0.50, *p =* .869, *d* = .08, 95% CI [.01, .15].

### Neuroimaging Results

The event-related functional analysis revealed 17 clusters that differentiated aware errors from unaware errors (Table 2). Aware errors were accompanied by greater activity in the left insula cortex (Figures 2C and 2D), the supramarginal gyrus (SMG; Figure 2B), and midline structures such as the left supplementary motor area (SMA), left anterior cingulate cortex (ACC) and bilateral precuneus (Figure 2A). It should be noted that while the centre of mass of activity in the SMA and ACC falls within the left hemisphere, the lateral extent of these clusters was bilateral.

**Figure 2.**
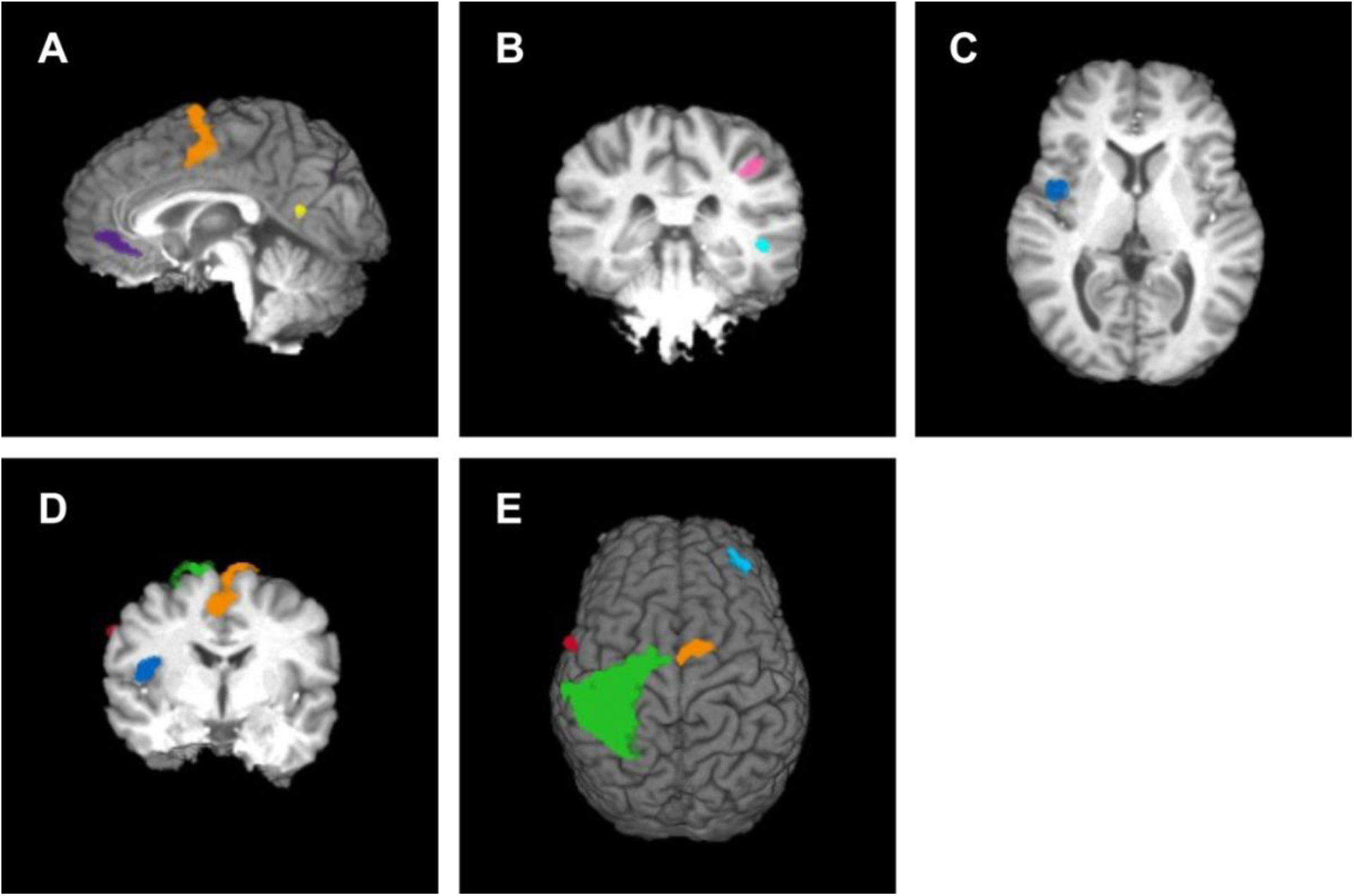
*Note*. Clusters associated with greater BOLD signal for aware errors than unaware errors. (A) Sagittal slice at *x* = 0. Purple cluster centred on left anterior cingulate cortex, orange cluster centred on left supplementary motor area (SMA), yellow cluster centred on left precuneus; (B) Coronal slice at *y* = 30. Blue cluster centred on right superior temporal gyrus, pink cluster centred on right supramarginal gyrus; (C) Axial slice at *z* = 10. Blue cluster centred on left insula; (D) Coronal slice at *y* = 5. Blue cluster centred on left insula, red cluster centred on left precentral gyrus, orange cluster centred on left SMA, green cluster centred on left postcentral gyrus; (E) Green cluster centred on right postcentral gyrus, orange cluster centred on left SMA, red cluster centred on left precentral gyrus, blue cluster centred on right middle frontal gyrus.

ACC activity was not robustly correlated with behavioural measures that are typically related to error awareness. That is, the speed of error commission was not significantly associated with the degree of ACC activity for either aware, *r*_*s*_ = -.09, *p* = .061, or unaware errors, *r*_*s*_ = -.06, *p* = .194. Further, we did not find evidence in support of an association between inhibition performance and ACC activity related to aware errors, *r*_*s*_ *=* .01, *p* = .890, nor error awareness rate, *r*_*s*_ *=* .09, *p* = .063. Likewise, we found no evidence for an association between post-error reaction time adjustments and ACC activity for aware errors, *r*_*s*_ = -.06, *p* = .233, or unaware errors, *r*_*s*_ = -.08, *p =* .096. The difference between BOLD activity in the ACC associated with aware errors and that associated with unaware errors was also not found to be significantly related to the speed of aware errors, *r*_*s*_ = -.02, *p* = 1.00, and unaware errors, *r*_*s*_ = .01, *p* = 1.00, or post-error adjustments in reaction time following aware errors, *r*_*s*_ = -.03, *p* = 1.00, and unaware errors, *r*_*s*_ = .09, *p* = .284.

The association between neural activity and performance indices was further assessed. Inhibition performance was found to correlate positively with BOLD activity associated with aware errors in the insula, *r*_*s*_ *=* .12, *p* = .030, and right SMG, *r*_*s*_ *=* .25, *p* < .001. Only the left middle frontal gyrus, *r*_*s*_ *=* .17, *p* = .010, and the SMA, *r*_*s*_ *=* .15, *p* = .030, were found to correlate significantly with aware error reaction time. Post-aware reaction time adjustments were associated with activity in the SMA, *r*_*s*_ = .15, *p* = .030, right SMG, *r*_*s*_ = .19, *p* < .001, right superior temporal gyrus, *r*_*s*_ = .14, *p =* .040, and left and right middle frontal gyri, *r*_*s*_ *=* .17, *p* = .010, and *r*_*s*_ *=* .21, *p* < .001, respectively, such that greater activity in these regions correlated with slower reaction time on the post-error trial. For unaware errors, neither the speed of the erroneous response nor post-error reaction time adjustments were found to significantly correlate with BOLD activity in any of the 17 clusters.

### Lasso Regression

Lasso regression results for each model are summarised in Table 3. Error awareness was found to be predicted by impulsivity, namely motor and planning scores from the BIS-11 (Barratt and Patton 1983), and behavioural inhibition score from the BIS/BAS (Carver and White 1994). The most important predictors of insula activity were attention to detail and imagination scores from the AQ (Baron-Cohen et al. 2001), and HADS depression score (Zigmond and Snaith 1983). No variable was found to predict ACC activity, while all variables except attention switching from the AQ, and attention, self-concept and DSM attention from the CAARS (Conners 1998) were found to predict SMG activity.

Regarding the questionnaire measures, it is worth noting that only a very small fraction of participants reported clinically relevant scores (see Table 4A of the Supplementary Material for descriptive statistics). The largest psychopathological subsample were individuals scoring in the clinical range for HADS anxiety (*n* = 196). Therefore, we compared the subsample of individuals meeting the cut-off for clinical levels of anxiety with those who did not. Corroborating the results of the lasso regressions, no difference was found between the groups in mean ACC, *t*(379) = 0.14, *p* > .990, insula, *t*(379) = 0.78, *p* > .990, SMG, *t*(379) = 2.10, *p* = .147, and middle frontal gyrus activity, *t*(379) = 0.77, *p* < .990.

## Discussion

The current study aimed to establish the robustness of previous findings on the neural correlates of error awareness. Here, we have discerned greater aware-related activity in a network of regions including the insula cortex, anterior cingulate cortex (ACC), supplementary motor area (SMA), and supramarginal gyrus (SMG). Further, individual differences in error-related neural activity were found to be related to indices of task performance in a select few regions including the insula, SMA and SMG. Moreover, we found that certain measures of psychopathology – namely impulsiveness and depression – explained variance in aware-related activity in a subset of these regions.

Although the ACC has been implicated in several studies on performance monitoring, differentiation of activity in this region with error awareness has been largely unreported (but see Hester et al., 2012). Our study has shown greater ACC activation – across the dACC and rACC – for aware errors than unaware errors, suggesting a sensitivity of the ACC to awareness. This supports claims that insufficient statistical power may underlie the discrepancy in previous findings (Wessel, 2012). Although the precise role of the ACC in error processing is unknown, there is a general consensus that the ACC – particularly the dACC – monitors ongoing behaviour and navigates the selection and evaluation of goal-directed behaviours (Holroyd and Yeung 2012). In particular, it is purported to respond to outcomes that are worse than expected and may signal the need for an adjustment in strategy to reach the desired goal (Holroyd and Coles 2002; Bryden et al. 2011). Our finding of greater rACC activation is not typically reported in error awareness studies, however the rACC has been proposed to be a neuronal generator of the Pe – an event-related potential associated with error awareness (Herrmann et al. 2004; Van Boxtel et al. 2005). While the rACC may be differentially involved in post-error processing, as evidenced by the Pe, further work is ultimately needed to discern precisely how the rACC contributes to awareness. Taken together, it is plausible that the ACC presents a threshold-like relationship to awareness and post-error processes, whereby a certain level of activity is sufficient to elicit error detection and post-error adaptation, but the overall level of activity is not tightly coupled to these processes (Orr and Hester 2012). This is moreover consistent with the absence of a relationship between individual differences in aware-related ACC activity and behavioural adjustments in our study. Thus, ACC activity may covary with error commission and contribute to error awareness such that it is facilitating goal attainment, however may not be solely responsible for eliciting awareness and post-error alterations.

Contrastingly, the insula cortex appears to be consistently modulated by error awareness. The insula has been proposed to be engaged in a number of processes, however its role in interoceptive awareness has taken prominence in recent decades. In particular, the insula integrates autonomic information with salient events such as errors (Klein et al. 2013b). Insula activity during aware errors may therefore be explained by interoceptive awareness of greater autonomic responses to aware errors (Craig 2009). Interestingly, concurrent insula and ACC activity during performance monitoring is a robust finding (Craig 2009; Ham et al. 2013). Although these structures are distinct, they have been purported to form a salience network which has been associated with interoceptive autonomic domains and the control of goal-directed behaviours (Dosenbach et al. 2007). Indeed, previous studies have found the ACC to be associated with autonomic engagement during aware error processing (Harsay et al. 2018). The relationship between the ACC and insula may explain how the ACC potentially mediates error awareness. Specifically, insula activity may represent awareness while ACC activity represents the control of directed effort. That is, error-related activity in the ACC may feedforward into the insula which may be more directly responsible for error awareness.

Consistent with previous findings, aware errors were associated with greater activity in the right SMG. The SMG is purported to be connected to the ACC and middle frontal cortex via the dorsal branch of the superior longitudinal fasciculus (SLF1; Ramos-Fresnedo et al. 2019). Recent research has demonstrated that individual differences in the SLF1 underpin an individual’s evidence accumulation capacity (Brosnan et al. 2020). This is pertinent given that current views on error awareness operate in line with an evidence accumulation account (Ullsperger et al. 2010). The emergence of error awareness is said to coincide with the accumulation of evidence above a response criterion threshold (Murphy et al. 2012). Given the SMG, ACC and middle frontal cortex were found to contribute to error awareness in the current study, it is plausible that connectivity between these regions might be a critical determinant of an individual’s error awareness.

It is also worth considering that the inferior parietal lobe – which in part comprises the SMG – has been proposed to form a network with the ACC and insula and together are associated with the salience of an event (Harsay et al., 2012). The parietal lobe, in particular, is suggested to act on salient events and likely works to direct and maintain the location of attention (Corbetta and Shulman 2002). Errors are arguably salient as they are infrequent and useful in that they re-direct a participant’s attention to current task goals. Indeed, consistent with this orienting account, we found elevated SMG activity to be correlated with slower reaction times following aware errors. This finding aligns with previous work which has found that correct trials following an error show heightened activation of the inferior parietal lobe, coinciding with increased post-error slowing (Marco-Pallarés et al. 2008).

Although error awareness rate did not appear to be associated with inhibition performance, we found a relationship between inhibition and aware-related activity in the insula and SMG. This is interesting given that error-related activity in the insula and inferior parietal lobe have previously been found to predict successful inhibition on the following No-Go trial (Hester et al. 2009b), suggesting a shared neural system between error awareness and successful response inhibition. Previously, we speculated that inclusion of two inhibition contingencies might disrupt the relationship between error awareness and future performance, reflecting the role of the ACC as a reinforcement learning signal (Orr and Hester 2012). We therefore opted to include only one inhibition rule in the current design. Despite this task change, we found no relationship between error awareness and inhibition performance. It is thus plausible that error awareness facilitates performance only under context-specific conditions – where there is a more direct contingency between an error and future performance. Since there was no direct contingency here, with performance not influencing the sequence of trials that followed, it is likely any increases in conservatism of responding are loosely, if at all, reflective of sustained changes in performance strategy.

While we examined post-error reaction time adjustments, it is worth considering that the error awareness task is not optimised for this analysis. Specifically, participants are required to make an awareness button press on the first post-error trial. To minimise this confound, we excluded the first two post-error trials from our post-error slowing analysis, however we still found greater slowing following unaware errors. While studies on post-error slowing and error awareness have generated mixed evidence (van Gaal et al. 2009; Hewig et al. 2011; Endrass et al. 2012; Hoonakker et al. 2016), the finding of greater slowing following unaware errors appears to be exclusive to studies employing the error awareness task. Given that post-error reaction time did not return to baseline by the third post-error trial, it seems plausible that unaware errors are accompanied by the continued anticipation of an impending No-Go trial, resulting in slowed responses. Our finding of greater slowing following unaware errors is therefore likely to be a task-specific phenomenon rather than a reflection of deliberate post-error behavioural adjustments. To reconcile these findings, we require a task that obviates the need for an error awareness button press on the post-error trial and offers more events (i.e., aware and unaware errors) per individual.

To examine the influence of dimensional measures of psychopathology on error awareness and related neural activity in four selected clusters (ACC, insula, SMG and middle frontal gyrus), we ran a series of lasso regressions. The most robust positive predictors of error awareness were impulsivity-related measures, specifically motor and non-planning impulsiveness. This is consistent with the finding that disorders marked by deficits in impulsiveness, such as ADHD and substance use disorder, have been shown to have impaired error awareness (O’Connell et al. 2009; Charles et al. 2017). Error awareness was also found to be positively predicted by behavioural inhibition system score which reflects the motivation to avoid adverse outcomes and is purported to be predictive of affective and behavioural responses after incentives and threats (Johnson et al. 2003). Moreover, the relationship between aware-related insula and SMG activity was found to be most notably positively predicted by depressive symptoms. Although some studies have found a heightened Pe – an event-related potential which is suggested to index error awareness – to be related to depressive symptoms (Mies et al. 2011; Mueller et al. 2015), others have found no such relationship (Compton et al. 2008). It has been reported, however, that depressed individuals display greater activity in the insula in response to negative stimuli than healthy controls (Hamilton et al. 2012). The heightened sensitivity to failure and negative information which is proposed to underlie clinical levels of depression may in part explain why aware-related activity in these regions is related to depressive traits in a non-clinical sample.

Our event-related analysis of a large sample revealed a network of regions including the insula cortex, SMG, and midline structures such as the ACC and SMA that show greater BOLD signal change for aware errors compared to unaware errors. The most parsimonious account of error awareness is that it is likely the result of the accumulative efforts of these systems which may not all individually drive awareness.

## Open practices statement

Data are available upon reasonable request and all scripts required for the current results have been made publicly available online at the Open Science Framework (https://osf.io/hrba7/).

## Acknowledgements

This work was supported by a Project Grant from the National Health and Medical Research Council (NHMRC) of Australia to M.A.B and R.H (#1045354). M.A.B is supported by a Senior Research Fellowship (Level B) from the NHMRC (# 1154378). This work was also supported by a Marie Skłodowska-Curie Fellowship from the European Commission (AGEING PLASTICITY; grant number 844246) to M.B, and supported by the NIHR Oxford Health Biomedical Research Centre. The Wellcome Centre for Integrative Neuroimaging is supported by core funding from the Wellcome Trust (203139/Z/16/Z). For the purpose of open access, the author has applied a CC BY public copyright licence to any Author Accepted Manuscript version arising from this submission. We would like to thank Mr Cameron Patrick for his assistance with the Lasso regression.

